# Extensive Hidden Prophage Diversity in *Enterobacter* Species Reveals Host Specificity and Local Distribution

**DOI:** 10.1101/2025.08.19.671078

**Authors:** Danna Paola Bours-Lugo, Juan Manuel Hurtado-Ramírez, Armando Hernández-Mendoza, Ramón González-Conde, Adrian Ochoa-Leyva, Gamaliel López-Leal

## Abstract

Bacteriophages are key drivers of bacterial evolution, particularly through their integration as prophages within host genomes. However, the diversity and host specificity of prophages in clinically relevant pathogens such as *Enterobacter* species remain poorly characterized. In this study, we revealed the diversity of prophages, mapped their distribution, and explored their relationships with their bacterial hosts. We analyzed 3,661 prophage sequences identified from the genomes of 20 different Enterobacter species. This analysis uncovered an extensive hidden diversity, comprising 1,617 phage genera and 2,423 phage species—nearly 80% of which were singletons—highlighting an exceptionally rich prophage landscape. We found substantial variation in prophage species richness across host species and isolation sources, with *E. kobei* and environmental isolates exhibiting the highest richness. Prophage populations showed strong host specificity and limited cross-species transmission. Moreover, these prophages exhibited geographic structuring and significant congruence between host and prophage phylogenies, as well as with the ecological lifestyles of their bacterial hosts. Although some interspecies transmission events were detected, they were infrequent. Overall, this study provides new insights into the diversity of *Enterobacter* prophages and underscores their ecological and clinical relevance in shaping host adaptation and phage–host dynamics.

**IMPORTANCE:** *Enterobacter* species are emerging opportunistic pathogens increasingly implicated in hospital-acquired infections. Although prophages play a pivotal role in bacterial genome evolution and host adaptation, their diversity and distribution across *Enterobacter* species remain largely uncharacterized. In this study, we performed a large-scale genomic analysis of 3,661 prophages from 20 *Enterobacter* species and uncovered an extensive and previously hidden prophage diversity. Our analysis revealed significant differences in prophage species richness across both host species and isolation sources, with *E. kobei* and environmental isolates exhibiting the highest richness. Prophage populations were strongly structured by host species and geography, showing limited cross-species transmission and a high degree of congruence between phage and host phylogenies. These findings highlight the structured and lineage-specific nature of prophage populations in *Enterobacter* and provide valuable insights into phage–host coevolution, microbial biogeography, and the design of targeted phage therapy strategies.

## INTRODUCTION

Viruses are the most abundant biological entities on Earth (1). Within this vast virosphere, bacteriophages (phages) specialize in infecting prokaryotic microorganisms (2). These phages replicate via the lytic cycle—typical of virulent phages—or by integrating into the host genome as prophages, or as cytoplasmic plasmid-like forms. Once integrated, prophages replicate alongside the host chromosome and transmit vertically from the original infected cell to its progeny during cell division (3). The integration of prophages into bacterial chromosomes can significantly alter host phenotypes and expand metabolic capabilities (4, 5). Some studies have reported that prophages also encode genes linked to antibiotic resistance (ARG) and virulence (VG) (6–9),thereby conferring adaptive advantages to their bacterial hosts and promoting horizontal gene transfer among microbes (10). Additionally, prophages can block infection by other phages that use the same receptor, providing superinfection immunity (11, 12). Also, the presence of orthologous prophages enables recombination with virulent phages, that could potentially generating defective particles, active dormant prophages (13)—or occasionally, producing novel infective viral variants (14, 15).

The growing threat of antimicrobial resistance constitutes a global public health crisis, contributing to an estimated 700,000 annual deaths due to drug-resistant bacterial infections (16). In this context, phages—nature’s bacterial predators—have re-emerged as a promising strategy to treat multidrug-resistant infections. In this sense, researchers have reported successful phage therapy cases targeting pathogens such as *Staphylococcus aureus* (17), *Pseudomonas aeruginosa* (18), *Acinetobacter baumannii*, and others (19, 20). However, despite significant progress in recent years, phage therapy remains in the experimental stage (21).

Importantly, many relevant pathogens carry abundant prophages in their genomes (22). Advances in nucleotide sequencing and bioinformatics have revealed that mobile genetic elements, including prophages, constitute a substantial portion of bacterial genomes (23). For example, in previous studies, we mined 13,713 complete bacterial genomes and found that 75% harbored identifiable prophages, with multiple bacterial genera showing an average of more than five prophages per genome—and at least one genome harboring more than twenty (22). Moreover, species within the genera *Enterobacter*, *Acinetobacter*, and *Pseudomonas* show significantly higher prophage enrichment in pathogenic isolates compared to nonpathogenic counterparts (22). Therefore, understanding and characterizing prophage populations in relevant pathogens is crucial to elucidate phage–host dynamics (24).

Species within the genus *Enterobacter* occupy diverse ecological niches within the *Enterobacteriaceae* family, including clinical, environmental, and animal-associated habitats (25). Moreover, several *Enterobacter* species act as plant pathogens (26) and also belong to the human gut commensal microbiota (27). Some species—such as *E. cloacae*, *E. asburiae*, and *E. hormaechei*—are frequent causes of hospital-acquired infections in humans (28). Due to their clinical impact and rising antibiotic resistance, the World Health Organization has classified *Enterobacter spp*. as critical priority pathogens (29). Only in the America continent, nearly 30–50% of *Enterobacter* isolates exhibit resistance to third-generation cephalosporins—a proportion that continues to rise (29). Consequently, alternative strategies such as phage therapy may offer promising options against these superbugs.

To date, no comprehensive studies have focused on profiling prophage populations across the entire *Enterobacter* genus. Existing work has targeted only a few species— such as *E. cloacae* (7), *E. asburiae* (30), and *E. hormaechei* (31)—but these analyses involved limited genome samples. On the other hand, only 117 genomes of phages infecting *Enterobacter* species had been deposited in NCBI (accessed October 2024). However, from this perspective, scientists have preferentially sequenced more bacterial than viral genomes—at least in the context of *Enterobacter* species—thereby unintentionally capturing a significant portion of their phages. As such, exploring prophage diversity represents a valuable resource for understanding phage–host dynamics.

In this study, we investigate phage and prophage diversity across multiple *Enterobacter* species. We analyzed a total of 3,778 bacteriophage sequences using comparative genomics and phylogenomics to deliver a comprehensive characterization of phage diversity and elucidate viral–host relationships.

## MATERIALS AND METHODS

### Genomes used and prophage identification

To conduct our study, 757 complete bacterial genomes of the *Enterobacter* genus were downloaded from NCBI. These genomes were downloaded in October 2024 from the NCBI. The quality of the genomes, namely, completeness and contamination percentages, was determined using CheckM2 (32). Only complete and uncontaminated genomes were included (completeness ≥95% and contamination ≤5%), and are listed in Supplemental Material Table S1. Biosample information for all genomes was obtained using efetch form E-utilities (33). We collected the metadata for the following sections: isolation source, isolation site, host, environmental medium, and sample type. We then categorized the isolates as Human, Animal, Clinical (this category corresponds to host isolates coming from instrumentation and equipment), and, Environmental. When this information was not available, it was grouped in the N/A category. Prophage predictions were carried out using VirSorter2 (34), and the quality of the prophage genomes was checked using CheckV (35). Only prophage sequences assigned as High-quality or Complete by checkv-quality were considered for downstream analysis. We followed the publicly available protocol for the validation of the first-instance prophage prediction (dx.doi.org/10.17504/protocols.io.bwm5pc86). Briefly, the final quality prophages analyzed by CheckV were validated in a second screening using VirSorter2. All genomes were annotated using prokka (36) and pharokka (37) for bacterial and viral genomes, respectively.

### Phylogenetic reconstruction of the genus *Enterobacter*

We aimed to construct homologous groups from *Enterobacter* bacterial genomes to gain insights into their evolutionary relationships. First, we ran Roary (38), setting the BLAST search parameters to a length coverage of ≥80% and an amino acid sequence identity of ≥80%. We created homologous groups (HG) with only one copy gene per genome, which we referred to as single-gene families (SGFs). SGFs were considered for the phylogenetic analysis. The SGFs were aligned with MUSCLE (39), specifying 50 iterations. Then, to create a DNA alignment in frame, we used the program TRANALING (40), which is part of the EMBOSS suite. Additionally, we discarded SGFs with recombination signals using PhiPack (41) with a *P*-value cutoff of 0.05. Then, the SGFs that did not show recombination signals were concatenated to form a super-alignment. Based on this alignment, we constructed a maximum likelihood (ML) phylogeny, selecting (TIM+F+I+G4) as the best model suggested by IQTree 2 (42). We ran a nonparametric bootstrap analysis (100 replicates) on the ML-phylogeny to establish the support for the clades. *Lelliottia nimipressuralis* (GCA_013898335) and *Lelliottia amnigena* (GCA_000016325) were used to give directionality to the tree (43).

### Bacteriophage clustering at the genus and species levels and phylogenetic reconstruction

To determine bacteriophage diversity, we first defined our prophage populations into species and genera following ICTV standards using vClust (44). Briefly, we performed an analysis similar to VIRIDIC (45) by calculating the total average nucleotide identity (tANI) between prophage genomes. Based on the total Average Nucleotide Identity (tANI) values, we classified viruses into species (≥95% tANI) and genera (≥70% tANI). We then constructed a Neighbor-Joining tree using the R ape (v5.6.1) and ggtree (v3.0.4) packages based on the pairwise identity matrix using only prophages that clustered into a genus with more than ≥5 members (to build a more robust and decisive tree). Similar, the prophage species with ≥7 members were visualized as an tANI (≥95%) network using Cytoscape v.3.8 (46). Bacteriophage taxonomic classification was carried out using taxMyPhage (47).

### Congruence between phylogenetic trees

To assess cophylogenetic patterns between *Enterobacter* hosts and their prophages, we conducted two complementary global-fit analyses: PACo (Procrustean Approach to Cophylogeny) (48) and ParaFit (49). First, we inferred host and prophage phylogenies independently (see above). These phylogenies were converted into pairwise patristic distance matrices and the host–phage associations were encoded as a binary matrix. The PACo analysis was performed using the paco package in R (v 0.4.2). The model was fitted using the default principal coordinate decomposition of distance matrices and Procrustean superimposition. Statistical significance of the global fit was evaluated using 10,000 permutations. ParaFit analysis was carried out using the vegan package in R (v2.6.4), with significance assessed via 999 permutations of the host–phage association matrix. To examine individual associations under the PACo framework, we computed squared residuals from the Procrustes superimposition. Associations with residual values above the 95th percentile threshold were classified as potential host-switching events, while associations below this cutoff were considered consistent with codivergence.

### Prophage diversity and geographical distribution

We evaluated the relationship between phage species or genera and both geographic origin, isolation source, and, host specie. To do this, we calculated the uncertainty coefficient (using the R DescTools v0.99.44) between prophage (phage genera and species levels) and host features such as geographic locality, isolation source and host species (*Enterobacter* species).

### Rarefaction Analysis of Prophage Species Richness

To compare prophage species richness while controlling for differences in sample depth, we performed rarefaction analyses using the vegan package in R (v2.6.4). Community matrices were constructed by aggregating prophage species counts per *Enterobacter* host species and per isolation source. For isolation sources, we included all categories and rarefied richness to 102 sequences, corresponding to 75% of the total sequences in the least-sampled source (Clinical). For host species, only those with ≥50 total sequences (prophages) were retained, and richness was rarefied to 63 sequences, corresponding to 75% of the total sequences in the least-sampled host (*E. ludwigii*). Using 75% of the total sequences in the least-sampled group as the rarefaction depth ensured that all categories were included in the analysis while minimizing the loss of diversity information from more deeply sampled groups. Rarefaction curves were generated using a fixed subsampling depth for all groups, and normalized richness values were extracted from the curves to enable direct comparison across categories.

## RESULTS

### High Abundance of Prophage Across *Enterobacter* species

To explore prophage populations across different *Enterobacter* species, we analyzed only high-quality and complete genomes (see Materials and Methods). We kept 747 genomes representing 20 distinct species, according to NCBI’s taxonomic classification (Supplementary Material: Table S1). However, to achieve more accurate species identification—given the complex taxonomy of *Enterobacter* species—we reconstructed a ML-phylogeny using 24 SGFs, and without recombination signals (see Materials and Methods). We first observed that most NCBI-assigned species clustered into well-supported clades (≥95% bootstrap support). Nevertheless, we reassigned 20.74% (155 genomes) of the genomes based on the ML-phylogeny (Figure 1A). For example, all genomes previously labeled by NCBI as *Enterobacter* spp. (Figure 1A: black labels), *E. roggenkampii* (orange labels), and *E. cloacae* (grass green labels) grouped within a major, well-supported clade corresponding to *E. hormaechei* (Figure 1A: blue labels). We therefore considered these genomes as *E. hormaechei* for all downstream analyses, and so on for all observed cases (Supplementary Material: Table S2). Importantly, the ML-phylogeny was consistent with previous studies (50). Finally, the most abundant species in our dataset were *E. hormaechei* (59.70%), *E. asburiae* (9.90%), *E. roggenkampii* (8.29%), *E. ludwigii* (5.48%), *E. kobei* (4.55%), and *E. cloacae* (4.28%). Interestingly, we identified a total of 3,661 high-quality prophages (see Materials and Methods), with a mode and mean of 5 and 4.9 prophages per genome, respectively. Prophages were present in all species analyzed, with 98.92% of genomes harboring at least one prophage. Remarkably, 39.49% of genomes contained between four and five prophages (Figure 1B). We then assessed whether prophage abundance was associated with the isolation source and/or *Enterobacter* species. Genomes isolated from human sources exhibited a significantly higher prophage in contrast with other sources (Figure 1C). Additionally, genomes belonging to *E. hormaechei* (*P*-value ≤ 0.01) and *E. asburiae* (*P*-value ≤ 0.05) were also enriched in prophages (Figure 1D). However, as noted above, more than 50% of our dataset corresponds to *E. hormaechei*, which could bias the statistical outcomes. To address this, we first assessed whether there were overall differences in prophage abundance among *Enterobacter* species. We applied a Kruskal–Wallis test (H = 89.44, p < 0.0001), which indicated that at least one species differed significantly from another in prophage abundance, meaning that prophage numbers were not homogeneous across species. We then conducted pairwise comparisons using two-tailed Mann–Whitney U tests with Holm–Bonferroni correction for multiple testing. The most significant differences involved *E. asburiae*, *E. bugandensis*, *E. ludwigii*, and *E. hormaechei* (Supplementary Material: Figure S1).

**Figure 1.**
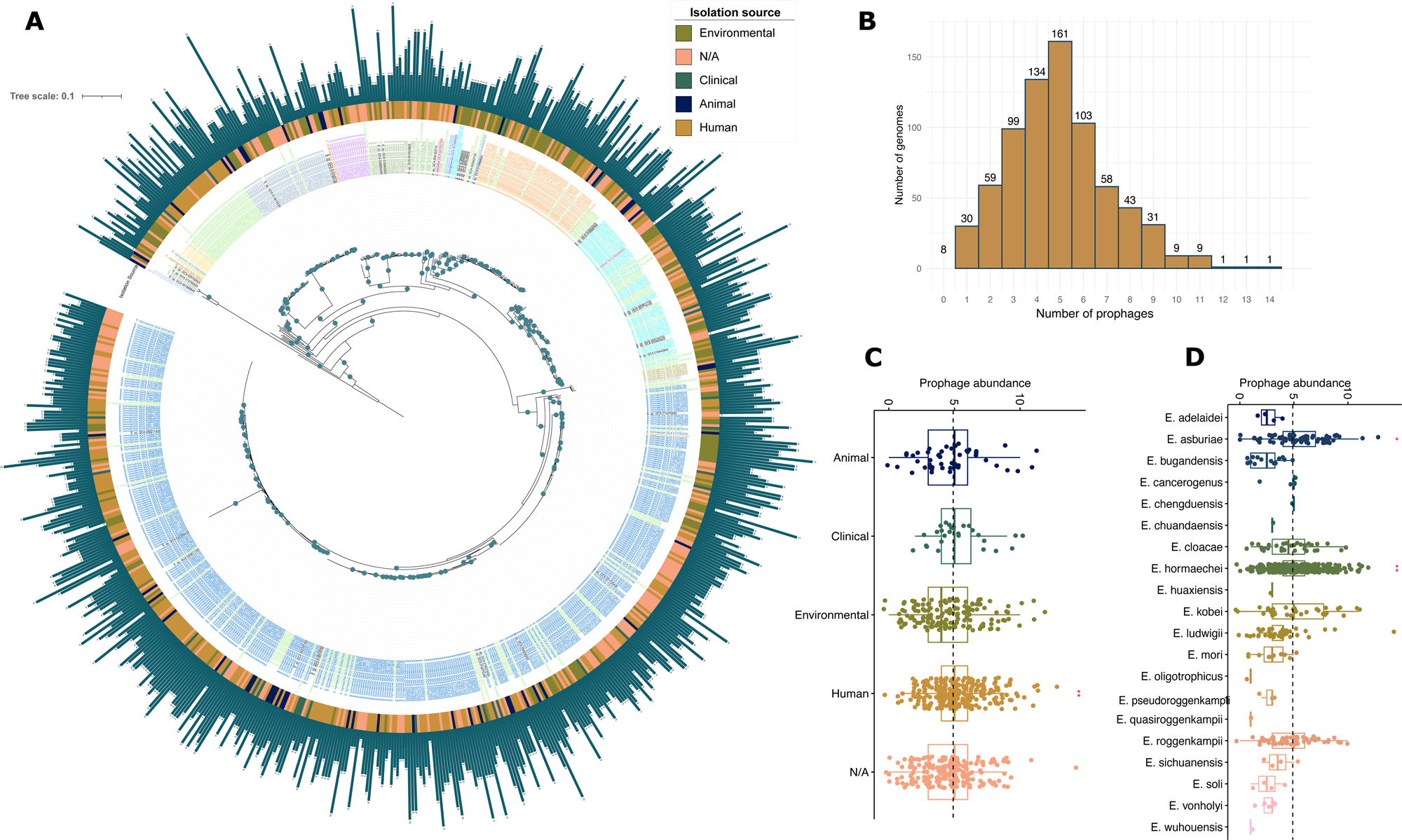
*Enterobacter* species phylogeny and prophage distribution. (A) Maximum likelihood (ML) phylogeny shows the relationships among all the genome. The external circle provides the isolation source of the isolates when available. Bar plots next to the isolation source circle denote the number of prophages identified for each genome. The colors of the labels denote the *Enterobacter* species, namely, *E. adelaidei* (#ff0000;red), *E. asburiae* (#39f9e8;turquoise), *E. bugandensis* (#bf7ff4;orchid), *E. cancerogenus* (#9e9f3d;yellowgreen), *E. chengduensis* (#1d36f4;blue), *E. chuandaensis* (#f28c04;darkorange), *E. cloacae complex*, *E. cloacae complex sp* and *E. cloacae* (#a1ea81;lightgreen), *E. dykesii* (#ffff2c;yellow), *E. hormaechei* (#076fcd;royalblue), *E. huaxiensis* (#d9ead3;gainsboro), *E. kobei* (#8eb27d;darkseagreen), *E. ludwigii* (#87a6c3;darkgray), *E. mori* (#ffd966;khaki), E. oligotrophicus (#bf9000;darkgoldenrod), *E. pasteurii* (#b4a7d6;lightsteelblue), *E. pseudoroggenkampii* (#15f6f4;aqua), *E. quasiroggenkampii* (#3d85c6;steelblue), *E. roggenkampii* (#f8a54d;sandybrown), *E. sichuanensis* (#49acf6;cornflowerblue), *E. soli* (#e06666;indianred), *E. sp*. (#000000;black), *E. vonholyi* (#e0cf9c;burlywood), *E. wuhouensis* (#7e9574;gray). The species *Lelliottia amnigena* (GCA_000016325) and *Lelliottia nimipressuralis* (GCA_013898335), shown in lightsteelblue, were used as an outgroup to root the phylogeny. The tree scale is the number of substitutions per site, and bootstrap values higher or equal to 95 are depicted with blue circles at the internal nodes of the phylogeny. (B) Bar graph depicting the distribution of total prophages in *Enterobacter* genomes. (C) Box plot representations of prophage abundance profiles according to host isolation source and Enterobacter species. Red asterisks indicate significant differences based on the Wilcoxon test (p-value ≤ 0.05; ** p-value ≤ 0.01).

### Extensive Hidden Prophage Diversity in *Enterobacter* species

To explore the diversity of *Enterobacter* prophages, we first contextualized our dataset by comparing the prophages identified in this study with 117 previously reported *Enterobacter* phage genomes retrieved from NCBI. We then defined phage species and genera by calculating pairwise genomic similarities among all sequences (see Materials and Methods). Interestingly, we identified 2,423 phage species, of which 79.98% were singletons (1,938 prophages), indicating these phage species were unique to individual genomes. *E. hormaechei* and *E. asburiae* were the species with the highest number of singleton prophages. However, given the uneven representation of *Enterobacter* species in our dataset, we normalized the number of singletons per species by dividing the number of singletons by the number of genomes analyzed for that species. *E. kobei* exhibited the highest singleton density (4.47 singletons per genome), although most species displayed densities ranging between 3 and 4 (Supplementary Material: Figure S2).

At the genus level, we identified 1,617 phage genera. Only genera comprising five or more members were used to construct a Neighbor-Joining phylogeny (Figure 2A). This tree revealed a high degree of genetic diversity among prophages, with many genera represented by only a few members. The most prevalent phage genera were Cluster 0, Cluster 1, and Cluster 2, with 57, 49, and 43 phages, respectively. Remarkably, members within these genera were nearly identical, as reflected by their extremely short branch lengths (Figure 2A; red branches). With the exception of the members of Cluster 0, which were found exclusively in *E. hormaechei*, the prophages of other major clusters (Clusters 1 and 2) were distributed across multiple host species. These highly conserved prophages were also found in genomes from different countries and isolation sources. In other words, only a few cases of cross-species transmission events were detected. Notably, prophages belonging to Cluster 1 were identified in six distinct *Enterobacter* species (Figure 2B). Additionally, prophages from Clusters 2, 3, 8, 35, 38, and 46 were also found in more than one *Enterobacter* species (Figure 2B; see dashed circles). However, these transmission events were less frequent compared to Cluster 1. Moreover, prophages from different isolation sources were also relatively rare (Figure 2B). In addition, prophage species occurring across different isolation sources were relatively uncommon. Most phage species were exclusive to a single isolation source, with only a few shared among different sources. The most notable overlap was observed between human-associated and environmental isolates, which shared 31 phage species in total (Supplementary Material: Figure S3).

**Figure 2.**
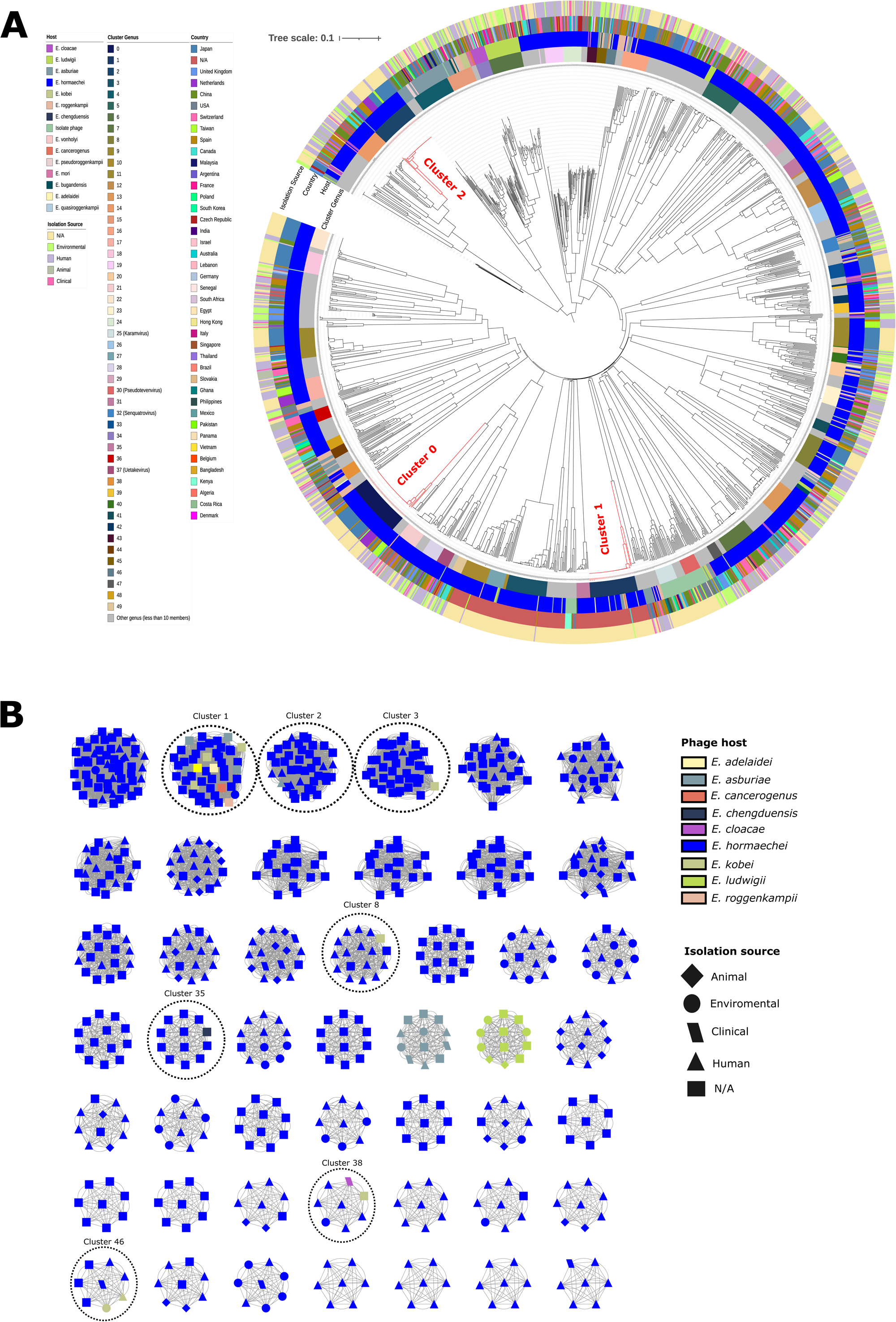
Phylogeny and Network Analysis of Enterobacter Phages. (A) The neighbor-joining (NJ) phylogenetic tree was constructed using 1,710 phage genomes, including only genera with more than five members, based on an intergenomic similarity matrix calculated with vClust at the genus level (≥70% tANI; see Materials and Methods). For clarity, only genera with ten or more members are shown in color (see cluster genus panel). The outer rings indicate the host isolation source, Enterobacter species, and country of origin. Clades corresponding to the three most prevalent genera are highlighted in red. (B) Network representation of *Enterobacter* bacteriophages at the species level, For the sake of clarity, only species having seven or more sequences are displayed (see Materials and Methods). Nodes are color-coded according to the *Enterobacter* species, and node shapes indicate the host isolation source. Dashed circles highlight the presence of prophages of the same species derived from different host species.

Although many clades included prophages from diverse geographic regions— suggesting broad global distribution—some clusters exhibited a moderate enrichment of prophages from specific countries, indicating possible regional circulation or localized transmission patterns. To evaluate whether prophage diversity is associated primarily with host species or geographic origin, we calculated the uncertainty coefficient (UC) between all phage species and genera with respect to host species and country of origin (Table 1). We found that the residual uncertainty was 90.94% considering phage species after knowing the country, and 75.83% for the phage genera. Conversely, the residual uncertainty for country after knowing the phage genus and species was 29.97% and 32.77%, respectively (Table 1). In other words, residual uncertainty determines how much information about a variable (X) you can obtain if you know another variable (Y), In this case, the UC is higher when species or genus is conditioned on country than the other way around, suggesting that geographic origin has a notable influence on phage linage at both genus and species levels. Furthermore, phage species provide more information about geographic origin than phage genera, and vice versa.

**Table 1.** Association between phage abundance and geographic or host-related variables based on the uncertainty coefficient. This table presents mutual information and conditional entropy values comparing bacteriophage genera and species with three host-related features: geographic origin (country), *Enterobacter* species, and isolation source. Mutual information quantifies the shared information between variables. The columns labeled “U(X|Y)” and “U(Y|X)” show the uncertainty coefficient, which indicates the proportion of uncertainty in variable X that is reduced by knowing Y (and vice versa), along with the corresponding confidence intervals (minimum “min” and maximum “max” values). Uncertainty coefficient values equal to or greater than 0.80 are highlighted in bold.

| X | Y | Mutual Information | U(X\|Y) | U(X\|Y)_min | U(X\|Y)_max | U(Y\|X) | U(Y\|X)_min | U(Y\|X)_max |
| --- | --- | --- | --- | --- | --- | --- | --- | --- |
| Species | Country | 2.428376 | **0.9094478** | **0.9031029** | **0.9157927** | 0.3277756 | 0.3225394 | 0.3330117 |
| Genus | Country | 2.025036 | 0.7583966 | 0.7487976 | 0.7679955 | 0.2997895 | 0.2940739 | 0.305505 |
| Species | Host | 1.437697 | **0.9208608** | **0.91344** | **0.9282817** | 0.1940572 | 0.1888739 | 0.1992405 |
| Genus | Host | 1.229766 | 0.7876813 | 0.7758634 | 0.7994992 | 0.1820571 | 0.1765355 | 0.1875787 |
| Species | Isolation source | 1.229766 | **0.8629423** | **0.853024** | **0.8728606** | 0.1556033 | 0.1528544 | 0.1583522 |
| Genus | Isolation source | 0.9056212 | 0.6779153 | 0.6642841 | 0.6915465 | 0.1340708 | 0.1309005 | 0.137241 |
| Host | Country | 0.3485117 | 0.1305227 | 0.1219662 | 0.1390792 | 0.2232282 | 0.2092817 | 0.2371746 |
| Host | Isolation source | 0.08563118 | 0.0641005 | 0.05511026 | 0.0730908 | 0.0548484 | 0.04753729 | 0.0621595 |

It is important to note that this analysis was initially performed on the complete dataset of 3,778 phages, suggesting that singleton sequences may have strongly influenced the results. However, when restricting the analysis to genera with more than 10 members, the UC decreased to 77.61% and 75.88% when evaluating phage species in relation to country and host species, respectively (Supplementary Material: Table S3). Next, when analyzing whether *Enterobacter* species were associated with geographic location and isolation source, we found that the uncertainty dropped to 13.05% and 9.41%, respectively (Supplementary Material: Table S3). These results indicate a clearer geographic structuring of phage populations, but not of *Enterobacter* species. Finally, we were able to assign a taxonomic classification at the genus level to only 3.46% of our dataset, using ICTV reference genomes (see Materials and Methods). Among these, only 36 prophages could be classified, with *Senquatrovirus* and *Novemvirus* being the most prevalent genera (Supplementary Material: Table S4). Moreover, all of them correspond to new phage species.

### Prophage species richness

We next assessed prophage species richness in relation to both bacterial host species and sources of isolation. Specifically, we quantified the diversity of prophages (prophage species) identified within each *Enterobacter* species, estimating how many distinct prophage species were associated with each host. Because most bacterial genomes in our dataset come from human-associated sources, and the majority were derived from *E. hormaechei*, we performed rarefaction analyses to compare prophage richness across *Enterobacter* host species and isolation sources, while controlling for differences in sampling effort.

For the analysis by isolation source, rarefaction was standardized to 102 sequences, representing 75% of the prophage count in the least sampled group (Clinical). Under this standardized effort, Environmental and Human-associated sources exhibited the highest expected richness (99.3 and 96.8 prophage species, respectively), while Animal and Clinical sources had lower values (90.4 and 89.3 prophage species, respectively; Figure 3A). Regarding host species, we included only those with at least 50 total prophage sequences and rarefied richness to 63 sequences, which corresponds to 75% of the prophage count from the least sampled host (*E. ludwigii*). Under these conditions, *E. kobei* displayed the highest expected richness (62.3 prophage species), followed by *E. cloacae* and *E. roggenkampii* (61.9 each), and then *E. asburiae* and *E. hormaechei* (60 each). E. ludwigii showed the lowest richness (52.9 prophage species) (Figure 3B).

**Figure 3.**
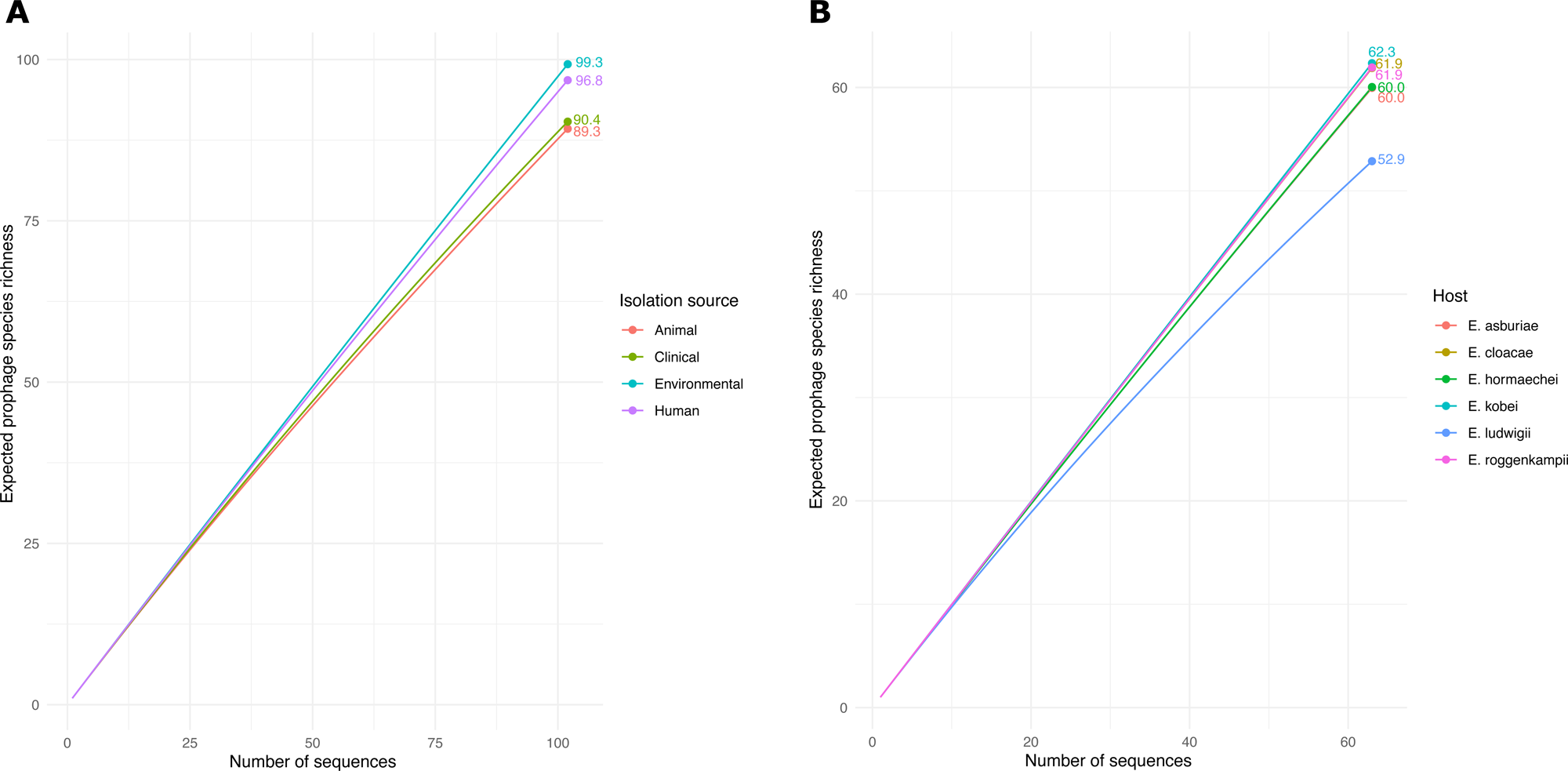
Rarefaction analysis of prophage species richness. Subsampling-based rarefaction curves were generated to estimate the expected richness of prophage species across different isolation sources (A) and *Enterobacter* species (B), while controlling for differences in sampling effort. Rarefaction for isolation sources was standardized, corresponding to 75% of the total sequences from the least sampled group (see Material and Methods).

Importantly, rarefaction curves revealed a rapid accumulation of novel prophage species with increasing sampling effort across both host species and isolation sources. These findings indicate that prophage diversity varies substantially depending on the bacterial host and ecological context, even after accounting for differences in sequencing depth. Notably, environmental and human-associated isolates, as well as certain Enterobacter species, harbor more diverse prophage populations.

### Coevolution and Host Switches Inferred by PACo Analysis

Cophylogeny arises when two sets of phylogenies and their interactions are congruent, which may indicate a shared evolutionary history (48). To assess the cophylogenetic signal between *Enterobacter* isolates and their prophages, we performed a PACo analysis using phylogenetic distances and host–phage association data (see Materials and Methods). The analysis revealed a significant global fit, resulting in a *P*-value of 0 after 10,000 permutations, indicating dependence of prophage phylogeny on host phylogeny. Additionally, the global test yielded a significant ParaFitGlobal (*P*-value of 0.001, 999 permutations), confirming a strong overall cophylogenetic signal between *Enterobacter* hosts and their prophages. These results are consistent with the observed host specificity in most prophage clusters and the limited number of cross-species transmission events (Figure 2B). We further investigated individual host–phage associations by calculating squared residuals, which quantify the deviation from the expected codivergence under the PACo model (1,650 associations). To identify potential host-switching events, we classified associations with residuals above the 95th percentile (Residual > 0.756) as candidate host jumps. Based on this threshold, we identified 83 potential host jumps (5.03%) and 1,567 associations consistent with codivergence (94.97%) (Figure 4).

**Figure 4.**
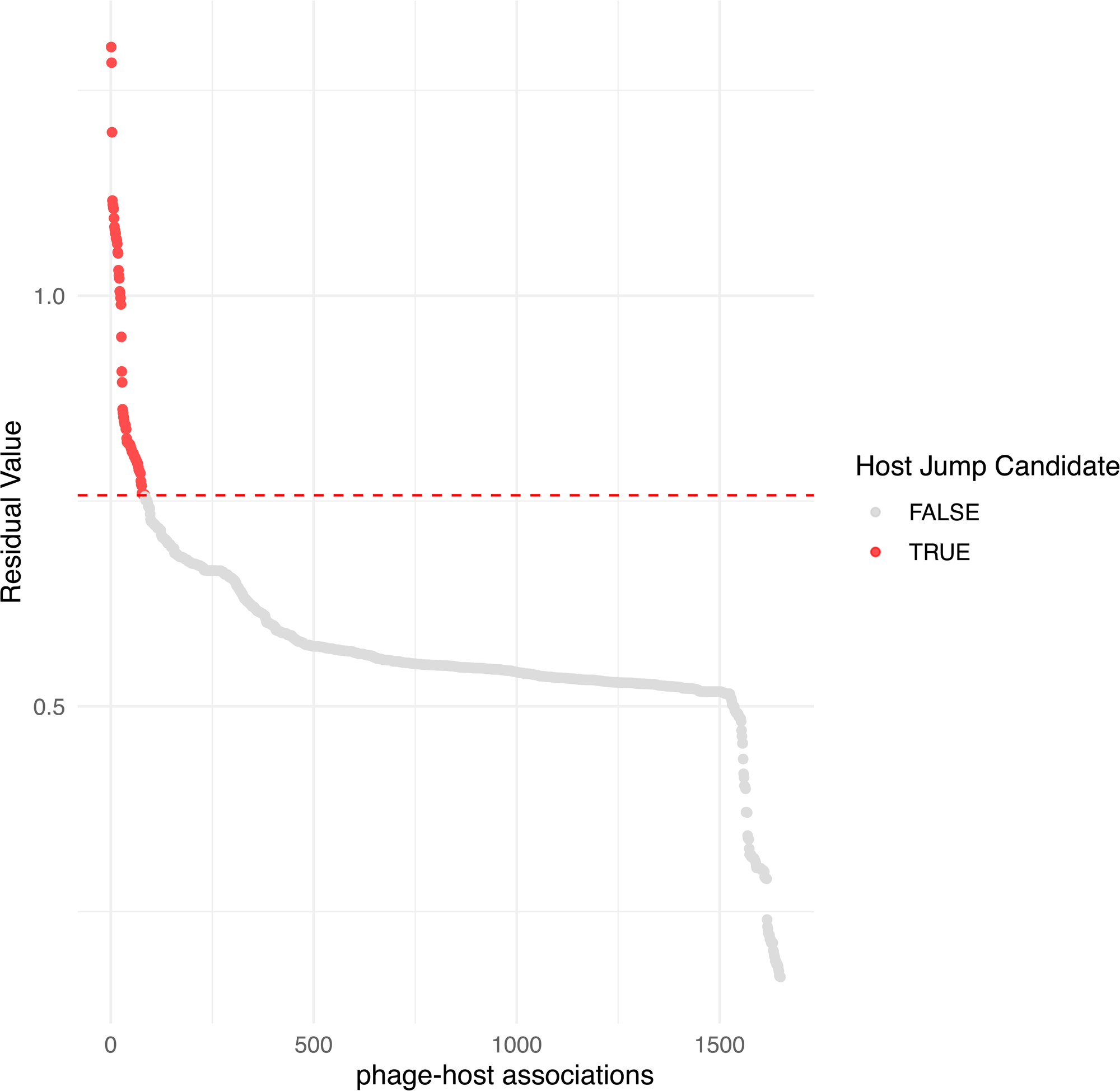
Plot of PACo residuals identifying candidate host jumps. Each point represents a unique association between an *Enterobacter* host and a prophage. Residual values (Y-axis) quantify the deviation of each association from the expected codivergence model, as inferred by PACo analysis (see Material and Methods). The X-axis ranks associations by increasing residual magnitude. Associations with residual values above the 95th percentile threshold (0.756; dashed red line) are highlighted in red and classified as potential host jump events (n = 83, 5.03%), while the remaining associations (n = 1,567, 94.97%) are consistent with codivergence.

## DISCUSSION

In recent years, phage therapy has re-emerged as a promising alternative to combat antibiotic resistance (21). In this context, the application of phage therapy in clinically relevant pathogens such as *E. cloacae*, *E. hormaechei*, *E. asburiae*, and *E. roggenkampii*—increasingly recognized as opportunistic pathogens of clinical concern— is considered a public health priority (7, 28–31). However, for such applications to be effective, it is critical to characterize the prophage populations in these species, as prophages can confer immunity to superinfection. Since integrated phage can block subsequent infection by related phages using the same bacterial receptor (11, 12).

A recent large-scale study analyzing over 13,000 bacterial genomes found that certain pathogens were enriched in prophage content. Notably, species within the genera *Acinetobacter*, *Pseudomonas*, and *Enterobacter* showed significant prophage enrichment (22). Another study analyzing complete genomes of 11,827 plasmids and 2,502 phages from the NCBI non-redundant database identified a large number of ARGs encoded within mobile elements (51). Additional studies focused specifically on *Enterobacter* species have also reported prophages carrying virulence or antibiotic resistance genes, although most of these analyses have been limited to a small number of *E. hormaechei* genomes associated with outbreaks (31). Although some work has investigated prophages in members of the *Enterobacteriaceae*, to our knowledge, no study has comprehensively characterized prophage populations across the genus *Enterobacter*. In this study, we analyzed 3,778 bacteriophage sequences derived from a curated set of high-quality, complete genomes representing diverse *Enterobacter* species. Importantly, these genomes were isolated from multiple sources, including humans, animals, and the environment. One of our main findings is the discovery of a previously unreported high prophage diversity. Specifically, we identified 1,617 phage genera and 2,423 species, of which only 25 genera were recognized by the ICTV. Additionally, 79.98% of prophages were singletons (1,938 in total), suggesting that many more phage genera and species remain to be discovered. A potential limitation of our study is the overrepresentation of *E. hormaechei* genomes, mostly derived from human sources. However, the density of singletons appeared comparable across several *Enterobacter* species. For reference, singleton frequencies in other pathogens such as *A. baumannii* have been reported to reach ∼70% (52, 53). These results reveal a vast diversity of prophages harbored by *Enterobacter* species, a pattern that also seems to occur in *Acinetobacter baumannii* (52, 53). In this context, we found that the diversity of prophage species remains largely unexplored. This suggests that broader and deeper sampling could uncover even greater prophage diversity. Notably, some human-associated pathogenic species such as *E. kobei* and *E. cloacae*, as well as *E. roggenkampii*—a pathogen reported in plants, animals, and humans—exhibited high prophage species diversity. Similarly, isolates from Environmental and Human-associated sources showed the highest expected prophage species richness. These results contribute to the definition of ‘viral dark matter’ (54), which refers to viral species that have not been characterized but whose existence has been revealed through metagenomic data and, in this case, by exploring host genomes. In contrast, prophage populations in other pathogens, such as *Streptococcus agalactiae* and *Campylobacter* species, appear to be less diverse (9, 55). Notably, there are no reports of inducible prophages in *S. agalactiae* or *Enterobacter* species, whereas such reports exist for *A. baumannii* (52). This highlights that the study of inducible prophages remains limited, as most research has focused on the identification of virulent phages.

From an ecological perspective, it is important to understand which phage species are circulating within specific geographic regions, hosts, and isolation sources. Interestingly, we found that phage species were more strongly associated with the country of origin, host species, and isolation source. Although this relationship was strongest when including singletons, the uncertainty coefficients remained high even when singletons were excluded from the analysis (see Materials and Methods). In other words, phages tend to be species-specific within *Enterobacter*. Furthermore, geography seems to influence the distribution of phage species, but not necessarily genera, suggesting local selective pressures or coevolutionary dynamics. In this sense, another important result is that host–prophage associations were not random and appear to be shaped by codiversification or host-specific ecological constraints. Our findings indicate that while most associations show strong phylogenetic congruence—suggesting long-term coevolution—there is also evidence of recent or past host-switching events involving a small subset of prophages, pointing to possible regional circulation or localized transmission patterns. Although this approach excludes a large proportion of singleton prophages, it ensures a more stable phylogenetic structure suitable for cophylogenetic analyses. This may bias the results toward more conserved or widespread phage lineages, but allows for statistically meaningful comparisons.

It should be noted that prophages from cluster 1 were found in six different species of *Enterobacter* from different countries, which was an atypical event. Exchange of prophages among isolates from different sources (e.g., environmental vs. clinical), even within the same host species, was rare, further indicating that prophage transmission may be constrained by ecological interactions or niche-specific adaptations. Collectively, these findings suggest that while prophages may occasionally cross host species boundaries, such events are infrequent and likely driven by specific biological or ecological circumstances. Therefore, circulating strains within distinct regions or ecological niches may influence susceptibility or resistance to certain phages.

From a public health perspective, it is also noteworthy that many of the ARGs identified in prophages were more frequently found in *Enterobacter* isolates from human sources. The rise in carbapenem-resistant *Enterobacter* species continues to be a global health concern (7, 51, 56). In this context, mobile genetic elements are recognized as key players in ARGs and VGs dissemination. While plasmids have been more extensively studied in this regard (51), recent evidence also implicates phages (31). In this context, we did not detect clear evidence of ARGs in our dataset using the Resistance Gene Identifier model (based on protein homology and SNPs). This absence may be attributed to our stringent criteria for selecting bona fide prophages, which ensured that only complete prophage genomes were retained. Although multiple studies have reported prophage-encoded ARGs, these are often found in degraded or incomplete prophages (57, 58).

Thus, considering all our results together, we propose two hypothetical scenarios. First, if these prophages become activated and propagate within host populations, they are likely to transfer such advantages (if they contain VGs or ARGs) only to very closely related hosts. This is probably due to phage-host coevolution, as supported by several lines of evidence. Geographic barriers and/or host lifestyle may further restrict this dissemination, this is likely because the majority of prophage species were associated with the isolation source of their respective bacterial hosts. In other words, prophage-mediated spread of ARGs or VGs across *Enterobacter* species is likely limited. Second, most *Enterobacter* prophages appear to be undergoing a process of domestication and may be doomed to persist passively within their host genomes, since domesticated prophages are under purify selection (13). Follow this idea, our results suggest that only “host jumps” candidates (see Figure 3) might remain active (5.03%). Although in this study we clearly found cases of interspecies transmission, further analyses combined with experimental evidence are needed to identify active prophages. From this perspective, the widespread presence of prophage signatures could serve more as a barrier against novel phage infections, where resident prophages promote the formation of defective particles (13).

Overall, our study provides valuable insights into the diversity, geographic distribution, and host dynamics of *Enterobacter* prophages. These findings underscore the importance of considering geographic origin and isolation source when designing phage therapy strategies—particularly against MDR pathogens such as *Enterobacter* species.

## ACKNOWLEDGMENTS

GLL thanks the Dirección General de Asuntos del Personal Académico (DGAPA) for awarding a postdoctoral fellowship for the period August 2015 to July 2016.

## FUNDING INFORMATION

This project was partially funded by CONAHCyT Ciencia Básica 2024 (grant no. CBF2023-2024-3171), given to GLL.

## SUPPLEMENTARY LEGENDS

**Supplementary Table S1.** List of bacterial genomes used.

**Supplementary Table S2.** List of bacterial genomes with new taxonomic assignation.

**Supplementary Table S3.** Relationship between phage species and geographic location assessed by the uncertainty coefficient.

**Supplementary Table S4.** List of bacteriophages taxonomically classified based on ICTV reference genomes using taxMyPhage (see Materials and Methods).

**Supplementary Figure S1.** Pairwise comparisons of prophage abundance across *Enterobacter* species. Pairwise statistical differences were evaluated using the Dunn test following a significant Kruskal–Wallis result (H = 89.44, p < 0.0001). The heatmap displays adjusted *p*-values from Holm-Bonferroni-corrected comparisons between *Enterobacter* species. Color intensity reflects statistical significance, with red indicating lower p-values (more significant differences) and blue indicating higher p-values (non-significant comparisons). Species are ordered alphabetically. Comparisons with adjusted p ≥ 0.05 are considered non-significant.

**Supplementary Figure S2.** Density of singleton prophage species per *Enterobacter* host. The barplot shows the density of singleton prophages identified in each *Enterobacter* species.

**Supplementary Figure S3.** Venn diagram of phage species found in *Enterobacter* genomes across different isolation sources. The diagram shows the overlap of prophage species identified in *Enterobacter* isolates from four distinct isolation sources: Clinical, Human, Environmental, and Animal (see Materials and Methods). Each area represents the set of unique phage species found in that category, with intersections indicating shared prophage species across sources.

## REFERENCES

1. Breitbart M, Rohwer F. 2005. Here a virus, there a virus, everywhere the same virus? Trends in Microbiology 13:278–284.

2. Clokie MR, Millard AD, Letarov AV, Heaphy S. 2011. Phages in nature. Bacteriophage 1:31–45.

3. Maurice CF, Bouvier C, de Wit R, Bouvier T. 2013. Linking the lytic and lysogenic bacteriophage cycles to environmental conditions, host physiology and their variability in coastal lagoons. Environmental Microbiology 15:2463–2475.

4. Ramisetty BCM, Sudhakari PA. 2019. Bacterial ‘Grounded’ Prophages: Hotspots for Genetic Renovation and Innovation. Frontiers in Genetics 10.

5. Suttle CA. 2007. Marine viruses--major players in the global ecosystem. Nat Rev Microbiol 5:801–12.

6. Costa AR, Monteiro R, Azeredo J. 2018. Genomic analysis of prophages reveals remarkable diversity and suggests profound impact on bacterial virulence and fitness. Scientific Reports 8.

7. Kondo K, Kawano M, Sugai M. 2021. Distribution of Antimicrobial Resistance and Virulence Genes within the Prophage-Associated Regions in Nosocomial Pathogens. Msphere 6.

8. López-Leal G, Santamaria RI, Cevallos MA, Gonzalez V, Castillo-Ramírez S. 2020. Prophages Encode Antibiotic Resistance Genes in. Microbial Drug Resistance 26:1275–1277.

9. Piña-González AM, Castelán-Sánchez HG, Hurtado-Ramírez JM, López-Leal G. 2024. prophage diversity reveals pervasive recombination between prophages from different species. Microbiology Spectrum 12.

10. Wendling CC, Refardt D, Hall AR. 2021. Fitness benefits to bacteria of carrying prophages and prophage-encoded antibiotic-resistance genes peak in different environments. Evolution 75:515–528.

11. Chung IY, Jang HJ, Bae HW, Cho YH. 2014. A phage protein that inhibits the bacterial ATPase required for type IV pilus assembly. Proceedings of the National Academy of Sciences of the United States of America 111:11503–11508.

12. Mcallister WT, Barrett CL. 1977. Superinfection Exclusion by Bacteriophage-T7. Journal of Virology 24:709–711.

13. Bobay LM, Touchon M, Rocha EPC. 2014. Pervasive domestication of defective prophages by bacteria. Proceedings of the National Academy of Sciences of the United States of America 111:12127–12132.

14. De Paepe M, Hutinet G, Son O, Amarir-Bouhram J, Schbath S, Petit MA. 2014. Temperate Phages Acquire DNA from Defective Prophages by Relaxed Homologous Recombination: The Role of Rad52-Like Recombinases. Plos Genetics 10.

15. Dragos A, Priyadarshini B, Hasan Z, Strube ML, Kempen PJ, Maróti G, Kaspar C, Bose B, Burton BM, Bischofs IB, Kovács AT. 2021. Pervasive prophage recombination occurs during evolution of spore-forming. Isme Journal 15:1344–1358.

16. Myers J. 2016. This is how many people antibiotic resistance could kill every year by 2050 if nothing is done. World Economic Forum.

17. Ramirez-Sanchez C, Gonzales F, Buckley M, Biswas B, Henry M, Deschenes MV, Horne B, Fackler J, Brownstein MJ, Schooley RT, Aslam S. 2021. Successful Treatment of Staphylococcus aureus Prosthetic Joint Infection with Bacteriophage Therapy. Viruses 13.

18. Onallah H, Hazan R, Nir-Paz R, group Ps, Brownstein MJ, Fackler JR, Horne B, Hopkins R, Basu S, Yerushalmy O, Alkalay-Oren S, Braunstein R, Rimon A, Gelman D, Khalifa L, Adler K, Abdalrhman M, Gelman S, Katvan E, Coppenhagen-Glazer S, Moses A, Oster Y, Dekel M, Ben-Ami R, Khoury A, Kedar DJ, Meijer SE, Ashkenazi I, Bishouty N, Yahav D, Shostak E, Livni G, Paul M, Gross M, Ormianer M, Aslam S, Ritter M, Urish KL, La Hoz RM, Khatami A, Britton PN, Lin RCY, Iredell JR, Petrovic-Fabijan A, Lynch S, Tamma PD, Yamshchikov A, Lesho E, Morales M, Werzen A, et al. 2023. Refractory Pseudomonas aeruginosa infections treated with phage PASA16: A compassionate use case series. Med 4:600–611 e4.

19. Schooley RT, Biswas B, Gill JJ, Hernandez-Morales A, Lancaster J, Lessor L, Barr JJ, Reed SL, Rohwer F, Benler S, Segall AM, Taplitz R, Smith DM, Kerr K, Kumaraswamy M, Nizet V, Lin L, McCauley MD, Strathdee SA, Benson CA, Pope RK, Leroux BM, Picel AC, Mateczun AJ, Cilwa KE, Regeimbal JM, Estrella LA, Wolfe DM, Henry MS, Quinones J, Salka S, Bishop-Lilly KA, Young R, Hamilton T. 2017. Development and Use of Personalized Bacteriophage-Based Therapeutic Cocktails To Treat a Patient with a Disseminated Resistant Infection. Antimicrobial Agents and Chemotherapy 61.

20. Wang ZT, Yang X, Wang H, Wang SX, Fang R, Li XT, Xing JY, Wu QQ, Li ZL, Song NN. 2024. Characterization and efficacy against carbapenem-resistant of a novel phage from sewage. Frontiers in Cellular and Infection Microbiology 14.

21. Segundo-Arizmendi N, Arellano-Maciel D, Rivera-Ramirez A, Pina-Gonzalez AM, Lopez-Leal G, Hernandez-Baltazar E. 2025. Bacteriophages: A Challenge for Antimicrobial Therapy. Microorganisms 13.

22. Lopez-Leal G, Camelo-Valera LC, Hurtado-Ramirez JM, Verleyen J, Castillo-Ramirez S, Reyes-Munoz A. 2022. Mining of Thousands of Prokaryotic Genomes Reveals High Abundance of Prophages with a Strictly Narrow Host Range. mSystems 7:e0032622.

23. Andrade-Martínez JS, Valera LCC, Cárdenas LAC, Forero-Junco L, López-Leal G, Moreno-Gallego JL, Rangel-Pineros G, Reyes A. 2022. Computational Tools for the Analysis of Uncultivated Phage Genomes. Microbiology and Molecular Biology Reviews 86.

24. Proenca M, Tanoeiro L, Fox JG, Vale FF. 2025. Prophage dynamics in gastric and enterohepatic environments: unraveling ecological barriers and adaptive transitions. ISME Commun 5:ycaf017.

25. Janda JM, Abbott SL. 2021. The Changing Face of the Family Enterobacteriaceae (Order: “Enterobacterales”): New Members, Taxonomic Issues, Geographic Expansion, and New Diseases and Disease Syndromes. Clin Microbiol Rev 34.

26. Garcia-Gonzalez T, Saenz-Hidalgo HK, Silva-Rojas HV, Morales-Nieto C, Vancheva T, Koebnik R, Avila-Quezada GD. 2018. Enterobacter cloacae, an Emerging Plant-Pathogenic Bacterium Affecting Chili Pepper Seedlings. Plant Pathol J 34:1–10.

27. Fei N, Zhao L. 2013. An opportunistic pathogen isolated from the gut of an obese human causes obesity in germfree mice. ISME J 7:880–4.

28. Davin-Regli A, Lavigne JP, Pages JM. 2019. Enterobacter spp.: Update on Taxonomy, Clinical Aspects, and Emerging Antimicrobial Resistance. Clin Microbiol Rev 32.

29. World Health O. 2017. WHO List of Critically Important Antimicrobials for Human Medicine (5th revision). World Health Organization, Geneva.

30. Islam MR, Mondol SM, Hossen MA, Khatun MP, Selim S, Amiruzzaman, Gomes DJ, Rahaman MM. 2025. First report on comprehensive genomic analysis of a multidrug-resistant Enterobacter asburiae isolated from diabetic foot infection from Bangladesh. Sci Rep 15:424.

31. Paauw A, Caspers MPM, Leverstein-van Hall MA, Schuren FHJ, Montijn RC, Verhoef J, Fluit AC. 2009. Identification of resistance and virulence factors in an epidemic Enterobacter hormaechei outbreak strain. Microbiology (Reading) 155:1478–1488.

32. Chklovski A, Parks DH, Woodcroft BJ, Tyson GW. 2023. CheckM2: a rapid, scalable and accurate tool for assessing microbial genome quality using machine learning. Nat Methods 20:1203–1212.

33. Sayers E. 2018. The E-utilities In-Depth: Parameters, Syntax and More. Accessed

34. Guo J, Bolduc B, Zayed AA, Varsani A, Dominguez-Huerta G, Delmont TO, Pratama AA, Gazitua MC, Vik D, Sullivan MB, Roux S. 2021. VirSorter2: a multi-classifier, expert-guided approach to detect diverse DNA and RNA viruses. Microbiome 9:37.

35. Nayfach S, Camargo AP, Schulz F, Eloe-Fadrosh E, Roux S, Kyrpides NC. 2021. CheckV assesses the quality and completeness of metagenome-assembled viral genomes. Nat Biotechnol 39:578–585.

36. Seemann T. 2014. Prokka: rapid prokaryotic genome annotation. Bioinformatics 30:2068–9.

37. Bouras G, Nepal R, Houtak G, Psaltis AJ, Wormald PJ, Vreugde S. 2023. Pharokka: a fast scalable bacteriophage annotation tool. Bioinformatics 39.

38. Page AJ, Cummins CA, Hunt M, Wong VK, Reuter S, Holden MT, Fookes M, Falush D, Keane JA, Parkhill J. 2015. Roary: rapid large-scale prokaryote pan genome analysis. Bioinformatics 31:3691–3.

39. Edgar RC. 2004. MUSCLE: a multiple sequence alignment method with reduced time and space complexity. BMC Bioinformatics 5:113.

40. Rice P, Longden I, Bleasby A. 2000. EMBOSS: the European Molecular Biology Open Software Suite. Trends Genet 16:276–7.

41. Bruen TC, Philippe H, Bryant D. 2006. A simple and robust statistical test for detecting the presence of recombination. Genetics 172:2665–81.

42. Minh BQ, Schmidt HA, Chernomor O, Schrempf D, Woodhams MD, von Haeseler A, Lanfear R. 2020. IQ-TREE 2: New Models and Efficient Methods for Phylogenetic Inference in the Genomic Era. Mol Biol Evol 37:1530–1534.

43. Alnajar S, Gupta RS. 2017. Phylogenomics and comparative genomic studies delineate six main clades within the family Enterobacteriaceae and support the reclassification of several polyphyletic members of the family. Infect Genet Evol 54:108–127.

44. Zielezinski A, Gudys A, Barylski J, Siminski K, Rozwalak P, Dutilh BE, Deorowicz S. 2025. Ultrafast and accurate sequence alignment and clustering of viral genomes. Nat Methods 22:1191–1194.

45. Moraru C, Varsani A, Kropinski AM. 2020. VIRIDIC-A Novel Tool to Calculate the Intergenomic Similarities of Prokaryote-Infecting Viruses. Viruses 12.

46. Otasek D, Morris JH, Boucas J, Pico AR, Demchak B. 2019. Cytoscape Automation: empowering workflow-based network analysis. Genome Biol 20:185.

47. Millard A, Denise R, Lestido M, Thomas MT, Webster D, Turner D, Sicheritz-Ponten T. 2025. taxMyPhage: Automated Taxonomy of dsDNA Phage Genomes at the Genus and Species Level. Phage (New Rochelle) 6:5–11.

48. Balbuena JA, Miguez-Lozano R, Blasco-Costa I. 2013. PACo: a novel procrustes application to cophylogenetic analysis. PLoS One 8:e61048.

49. Legendre P, Desdevises Y, Bazin E. 2002. A statistical test for host-parasite coevolution. Syst Biol 51:217–34.

50. Wu W, Feng Y, Zong Z. 2020. Precise Species Identification for Enterobacter: a Genome Sequence-Based Study with Reporting of Two Novel Species, Enterobacter quasiroggenkampii sp. nov. and Enterobacter quasimori sp. nov. mSystems 5.

51. Jiang L, Zhu H, Wei J, Jiang L, Li Y, Li R, Wang Z, Wang M. 2023. Enterobacteriaceae genome-wide analysis reveals roles for P1-like phage-plasmids in transmission of mcr-1, tetX4 and other antibiotic resistance genes. Genomics 115:110572.

52. Arellano-Maciel D, Hurtado-Ramirez JM, Camelo-Valera LC, Castillo-Ramirez S, Reyes A, Lopez-Leal G. 2025. Geographic variation in abundance and diversity of Acinetobacter baumannii Vieuvirus bacteriophages. Front Microbiol 16:1522711.

53. Tenorio-Carnalla K, Aguilar-Vera A, Hernandez-Alvarez AJ, Lopez-Leal G, Mateo-Estrada V, Santamaria RI, Castillo-Ramirez S. 2024. Host population structure and species resolution reveal prophage transmission dynamics. mBio doi:10.1128/mbio.02377-24:e0237724.

54. Chevallereau A, Pons BJ, van Houte S, Westra ER. 2022. Interactions between bacterial and phage communities in natural environments. Nature Reviews Microbiology 20:49–62.

55. Kovacec V, Di Gregorio S, Pajon M, Crestani C, Poklepovich T, Campos J, Basit Khan U, Bentley SD, Jamrozy D, Mollerach M, Bonofiglio L. 2024. Revisiting typing systems for group B Streptococcus prophages: an application in prophage detection and classification in group B Streptococcus isolates from Argentina. Microb Genom 10.

56. Rodriguez-Bano J, Gutierrez-Gutierrez B, Machuca I, Pascual A. 2018. Treatment of Infections Caused by Extended-Spectrum-Beta-Lactamase-, AmpC-, and Carbapenemase-Producing Enterobacteriaceae. Clin Microbiol Rev 31.

57. Kondo K, Kawano M, Sugai M. 2021. Distribution of Antimicrobial Resistance and Virulence Genes within the Prophage-Associated Regions in Nosocomial Pathogens. mSphere 6:e0045221.

58. Takeuchi N, Hamada-Zhu S, Suzuki H. 2023. Prophages and plasmids can display opposite trends in the types of accessory genes they carry. Proc Biol Sci 290:20231088.

